# Inactivation of LCN2/EGR1 Promotes Oligodendrocyte Progenitor Cell Differentiation and Remyelination after White Matter Injury

**DOI:** 10.1101/2020.01.02.892976

**Authors:** Qiang Li, Xufang Ru, Yang Yang, Hengli Zhao, Jie Qu, Weixiang Chen, Pengyu Pan, Huaizhen Ruan, Chaojun Li, Hua Feng, Yujie Chen

## Abstract

The insufficient remyelination due to the impaired oligodendrocyte precursor cell differentiation and maturation is highly associated with irreversible white matter injury and neurological deficits. Consequently, inhibitory components and microenvironment for remyelination might serve as potential therapeutic targets for treating white matter injury after acute central nervous system injury and neurodegeneration diseases. Lipocalin-2 was recently reported to corelate with white matter in both atypical, acute white matter injured disease subarachnoid hemorrhage and typical, chronic white matter injured disease multiple sclerosis. To elucidate the role and underlying mechanism of Lipocalin-2 in oligodendrocyte precursor cell differentiation and remyelination, we used genetic inhibition and a constitutive conditional knockout model with subarachnoid hemorrhage or multiple sclerosis. We found that the genetic inhibition of the increase in Lipocalin-2 promoted oligodendrocyte precursor cell differentiation, remyelination, and functional recovery after subarachnoid hemorrhage or multiple sclerosis. Unexpectedly, the inhibition of Lipocalin-2 did not reduce glial activation and inflammation. Lipocalin-2 was shown to activate Early Growth Response Protein 1 in oligodendrocyte precursor cells, which is partly regulated by its receptor SLC22A17. In the conditional knockout of Early Growth Response Protein 1 in oligodendrocyte precursor cells, we discovered enhanced oligodendrocyte precursor cell differentiation in developing and injured white matter; consistently, the specific inactivation of Early Growth Response Protein 1 promoted remyelination and neurological recovery after subarachnoid hemorrhage or multiple sclerosis. Thus, we propose that following white matter injury in humans, the increase in Lipocalin-2 activates Early Growth Response Protein 1 and consequently impair oligodendrocyte precursor cell differentiation and myelin repair. Our results suggest that therapies specifically inactivating Lipocalin-2/ Early Growth Response Protein 1 signal in oligodendroglial lineage cells could represent a novel strategy to enhance differentiation and remyelination in white matter injury patients.

## Introduction

White matter injury (WMI) causes deficits in movement, sensation and cognition and is observed in typical demyelination disorders such as multiple sclerosis (MS), amyotrophic lateral sclerosis and experimental autoimmune encephalomyelitis(van Tilborg et al, 2018). Similarly, in hemorrhagic stroke, including subarachnoid hemorrhage (SAH), WMI remains a pivotal factor that impedes functional recovery and unsatisfactory prognosis after surgical rescue(Egashira et al, 2015; Macdonald & Schweizer, 2017; Wu et al, 2017).

Although WMIs have distinct etiologies, a shared neuropathological feature involves the failed differentiation of oligodendrocyte precursor cells (OPCs) into myelin-producing oligodendrocytes following injury, causing impaired myelin formation and regeneration (remyelination). Nevertheless, the mechanisms underlying this pathology have not been fully elucidated, as there is a lack of comprehensive understanding of the inhibitory microenvironment that deters oligodendrocyte differentiation and regeneration; moreover, there is a lack of approved therapies(Kremer et al, 2011).

Previous work found that Lipocalin-2 (LCN2) plays a negative role in remyelination after inflammatory stimuli in the central nervous system and suggested that LCN2 is a promising remyelination therapeutic target(Al Nimer et al, 2016; Ngo et al, 2015). Moreover, these studies indicated that LCN2 activates glial cells, which then secretes proinflammatory cytokines to damage myelin(Nam et al, 2014). This interpretation may be not universal since LCN2 does not induce glial activation in some demyelinating diseases, such as Alzheimer’s disease(Dekens et al, 2018). Given that OPCs are involved in remyelination under many conditions, an investigation into whether LCN2 affects the function of OPCs after demyelination is warranted. Recent observations in a genetically deficient mouse model showed that LCN2 was associated with WMI after intracranial hemorrhage and SAH, although the specific mechanism was not elucidated(Egashira et al, 2014).

Moreover, we observed the remarkable proliferation and migration of OPC after SAH, but the remyelination was incomplete and insufficient to restore long-term neurological function, suggesting a potent inhibitory microenvironment that impedes oligodendrocyte differentiation and maturation following SAH and in other conditions(Li et al, 2018). In addition, LCN2 paucity reduced the expression of NG2 (an OPC marker) and ameliorated neurological deficits(Egashira et al, 2014). Hence, the questions of whether LCN2, as one member of the inhibitory microenvironment, inhibits remyelination by suppressing OPC differentiation and how to regulate oligodendroglial lineage cell behavior remain to be fully elucidated.

Here, we reveal the roles of LCN2 in the remyelination and functional recovery of WMI diseases, such as SAH and MS. Furthermore, we demonstrate the underlying mechanism of how LCN2 activation regulates oligodendrocyte lineage cell responses, revealing potentially targetable receptors for clinical intervention in myelin disorders throughout the lifespan.

## Results

### Enrichment of LCN2 impeded functional recovery after WMI

To detect whether LCN2 is a shared overexpressed pathological molecule after WMI, we sought to thoroughly evaluate LCN2 expression in typical white matter disorders (MS) and atypical white matter disorders (SAH). First, an endovascular perforation (E.P.) was used to induce SAH, and a cuprizone diet was used to induce a MS model (Fig. 1a, c). According to immunofluorescence analysis, LCN2 expression was enhanced in the brains (especially in the white matter area) of SAH mice and in the spinal cords of MS mice (Fig. 1b, d). Upregulation of LCN2 was confirmed not only in the acute phages (within 2 days) of SAH, as previously verified by an immunohistochemical method(Egashira et al, 2014), but also over a longer period (1-14 days) following SAH, as determined by western blot analysis (Online Resource Supplemental Fig. 1e).

**Fig. 1.**
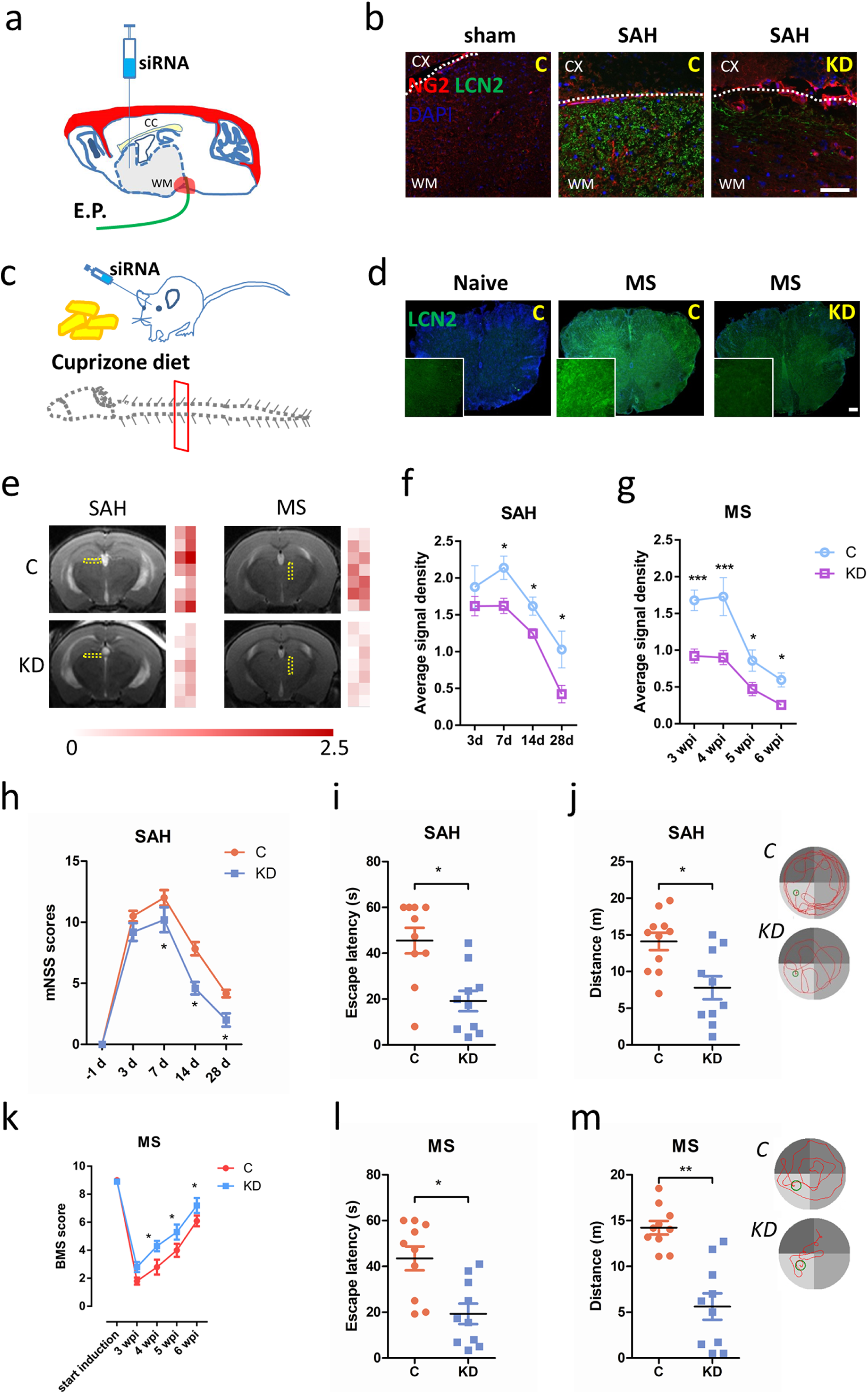
Enrichment of LCN2 impeded functional recovery after WMI. **a)** Schematic of the E.P.-induced SAH model and injection of siRNA. **b)** Image of NG2 (red) and LCN2 (green) expression in the brains of control and LCN2*^KD^* mice at 7 days after sham or SAH. Scale bar, 50 μm. **c)** Schematic of the cuprizone diet-induced MS model and injection of siRNA. **d)** Image of LCN2 (green) expression in the spinal cords of control and LCN2*^KD^* mice at 3 wpi of MS. Scale bar, 50 μm. **e)** MRI (T2WI) of control and LCN2*^KD^* mice at 7 days after SAH and at 3 wpi of MS, respectively. The colored grid bar on the right of each panel shows the T2WI signal of the yellow frame, ranging from 0 to 2.5. **f)** Average signal density of the white matter of control and LCN2*^KD^* mice (n = 3 per time point), at 3, 7, 14, and 28 days after SAH. Two-way ANOVA with Sidak’s multiple comparisons test, *P < 0.05. **g)** Average signal density of the spinal cord white matter of control and LCN2*^KD^* mice (n = 3 per time point) at 3, 4, 5, and 6 wpi of MS. Two-way ANOVA with Sidak’s multiple comparisons test, *P < 0.05, ***P < 0.001. **h)** mNSS scores of control and LCN2*^KD^* mice (n = 6 per time point) at -1 (1 day before SAH induction), 3, 7, 14, and 28 days after SAH. Two-way ANOVA with Sidak’s multiple comparisons test, *P < 0.05. **i)** Escape latency before finding the platform of control and LCN2*^KD^* mice (n = 10 each group) with SAH. Two-tailed Student’s t test, *P = 0.0342. **j)** Swimming distance before finding the platform of control and LCN2*^KD^* mice (n = 10 each group) with SAH. Two-tailed Student’s t test, *P = 0.0406. **k)** BMS scores of control and LCN2*^KD^* mice (n = 6 per time point) at 0 (induction), 3, 4, 5, and 6 wpi of MS. Two-way ANOVA with Sidak’s multiple comparisons test, *P < 0.05. **l)** Escape latency before finding the platform of control and LCN2*^KD^* mice (n = 10 each group) with SAH. Two-tailed Student’s t test, *P = 0.0212. m) Swimming distance before finding the platform of control and LCN2*^KD^* mice (n = 10 each group) with SAH. Two-tailed Student’s t test, **P = 0.0052.

Next, to further assess whether blocking this enrichment of LCN2 benefited white matter functional recovery, we established an effective knockdown in which LCN2 was obviously repressed due to siRNA injection (Fig. 1a, c). The knockdown of LCN2 (KD) significantly inhibited its enhancement in the white matter (Fig. 1b, d) and reduced LCN2 protein expression and concentration in the cortical cortex and corpus callosum, as determined by western blotting and ELISA (Online Resource Supplemental Fig. 1f, g).

Moreover, to verify whether the inhibition of LCN2 affected WMI, we detected the T2 signal of the brains of mice in the MS and SAH groups by 7.0T magnetic resonance (Fig. 1e). In the corpus callosum of SAH mice and in the white matter of MS mice, the average T2 signal density, which predicts brain injury, was significantly reduced after LCN2 knockdown (Fig. 1f). Moreover, WMI volume after SAH seemed to decrease in LCN2*^KD^* mice (Online Resource Supplemental Fig. 1c). Nonetheless, the knockdown of LCN2 did not affect rebleeding or hematoma absorption because the hemorrhagic severity of endovascular perforation induced SAH was not significantly changed by bleeding volume score and MRI (Online Resource Supplemental Fig. 1a, b). The secretion of LCN2 increased as the hemorrhagic score increased in the white matter at 3 days after SAH (Online Resource Supplemental Fig. 1d).

Finally, to determine whether the knockdown of LCN2 promoted functional recovery after WMI, we assayed motor, learning and memory functions in SAH and MS mice using neurological scoring and the Morris water maze. The knockdown of LCN2 reduced mNSS scores in SAH mice at 7, 14 and 28 days and improved BMS scores at 4, 5, and 6 weeks after MS (Fig. 1g, j). Furthermore, the knockdown of LCN2 significantly shortened the escape latency and searching distance before finding the underwater platform (Fig. 1h, i, k, l). Overall, these data suggest that the upregulation of LCN2 impedes functional recovery after WMI.

### Inhibition of LCN2 promotes remyelination and myelin maturation after WMI

Having shown the positive effects of LCN2 reticence on functional recovery after WMI, we next asked whether the inhibition of LCN2 could promote remyelination. Previous work showed that LCN2 contributes to demyelination in MS, spinal cord injury and optic neuritis, as well as SAH (Chun et al, 2015; Khalil et al, 2016; Rathore et al, 2011). We also found that the inhibition of LCN2 reduced the expression of degraded myelin basic protein (DMBP), a biomarker of demyelination, while regulating MBP in the white matter of LCN2*^KD^* mice after SAH (Online Resource Supplemental Fig. 2a, b). Nevertheless, the role of LCN2 in myelination is controversial because some studies indicate that it is neuroprotective(Ferreira et al, 2015). To our surprise, the double-labeling results showed that compared with scrambled control treatment, the knockdown of LCN2 significantly increased remyelination at 7, 14, and 28 days post-SAH, as measured by remyelination index (colocalization of MBP and axonal neurofilament 200 (NF200), divided by area of neurofilament) (Fig. 2a, b). Black Gold Ⅱ staining provided morphological confirmation, as LCN2*^KD^* mice showed more myelinated fibers and higher BG intensity than control mice(Fig. 2d-f). Similarly, in the MS model, an increase in the level of MAG, an oligodendrocyte maturation-associated gene, was observed in LCN2*^KD^* mice compared to control mice (Fig. 2a, c). Given that LCN2 was considered a promising prognostic biomarker for early MS patients(Khalil et al, 2016), we next used Black Gold Ⅱ staining to determine whether the knockdown of LCN2 also promoted remyelination after MS. The results indicated an increased number of myelinated fibers and therefore an increase in the remyelination index in the corpus callosum and white matter at 5 weeks after MS (Fig. 2g-i). To examine whether LCN2 knockdown affected the myelin sheath during the remyelination process, we analyzed the ultrastructure of axon and myelin integrity in the white matter. Our previous observation revealed SAH decreased myelin thickness in the corpus callosum and white matter during early phage(Li et al, 2018). At 7 days after SAH in LCN2*^KD^* mice, we detected increased myelin thickness and statistically significantly reduced mean g-ratio, a measure of myelin sheath thickness calculated as the ratio of axon diameter to myelinated fiber diameter, in LCN2*^KD^* mice compared to scrambled control-treated mice (Fig. 2j, k). At 5 weeks after MS, the mean g-ratio of the myelinated axons of LCN2*^KD^* mice was significantly lower than that of control mice, indicating that the myelin sheaths were thicker in LCN2*^KD^* mice (Fig. 2j, l).

**Fig. 2.**
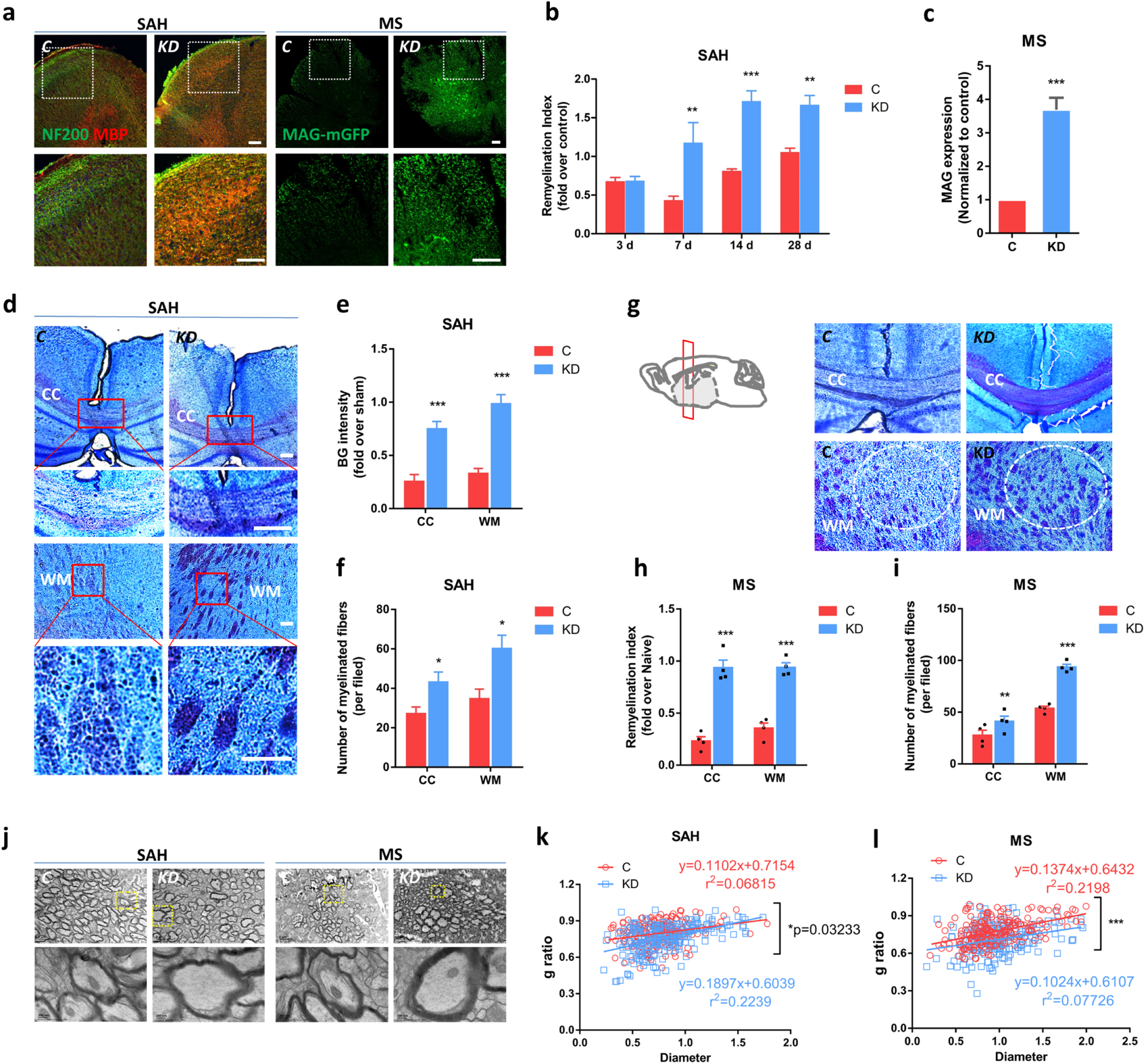
Inhibition of LCN2 promotes remyelination and myelin maturation after WMI. **a)** Image of NF200 and MBP in the white matter (WM) of control and LCN2*^KD^* mice at 7 days after SAH, with expanded field of white frame, respectively; Image of MAG (green) in the spinal cord WM of control and LCN2*^KD^* mice at 5 wpi of MS, with expanded field of white frame. Scale bar, 50 μm. **b)** Remyelination index (colocalization of MBP and NF200, normalized to area of NF200) of control and LCN2*^KD^* mice (n = 5 fields per time point) at 3, 7, 14, and 28 days after SAH. One-way ANOVA with Dunnett’s multiple comparisons test, **P < 0.01, ***P < 0.001 vs. the control group. **c)** Normalized MAG expression of control and LCN2*^KD^* mice at 5 wpi of MS (n = 5 fields per time point). Two-tailed Student’s t test, ***P = 0.0003. **d)** Black Gold (BG) II staining of the brains of control and LCN2*^KD^* mice at 7 days after SAH, with expanded fields of the corpus callosum (CC) and WM. Scale bar, 75 μm. **e)** Relative BG intensity in the CC and WM of control and LCN2*^KD^* mice (n = 5 fields per time point) at 7 days after SAH. One-way ANOVA with Dunnett’s’ multiple comparisons test, ***P < 0.001, vs. control group. **f)** Number of myelinated fibers in the CC and WM of control and LCN2*^KD^* mice (n = 5 fields per time point) at 7 days after SAH. One-way ANOVA with Dunnett’s multiple comparisons test, *P < 0.05, vs. the control group. **g)** BG II staining of the CC and WM of control and LCN2*^KD^* mice at 5 wpi inof MS. Scale bar, 75 μm. **h)** Normalized remyelination index in the CC and WM of control and LCN2*^KD^* mice (n = 5 fields per time point) at 5 wpi of MS. One-way ANOVA with Dunnett’s multiple comparisons test, ***P < 0.001, vs. the control group. **i)** Number of myelinated fibers in the CC and WM of control and LCN2*^KD^* mice (n = 5 fields per time point) at 5 wpi of MS. One-way ANOVA with Dunnett’s’ multiple comparisons test, **P < 0.01, ***P < 0.001, vs. the control group. **j)** Ultrastructure of the WM of control and LCN2*^KD^* mice at 7 days after SAH and at 5 wpi at MS. The yellow frames are magnified below. Scale bar, 2 μm or 200 nm. **k)** G-ratio versus axon diameter in control and LCN2*^KD^* mice at 7 days after SAH. Extra sum of squares F test between slopes, * P = 0.00323. **l)** ratio versus axon diameter in control and LCN2*^KD^* mice at 5 wpi at MS. Extra sum of squares F test between slopes, ***P = 0.0001.

We further investigated whether LCN2 regulates myelin membrane compaction/maturation. During the remyelination phages of SAH or MS, the prominent enlargement of the inner myelin tongues of LCN2*^KD^* mice was associated with small-diameter axons (0.4<diameter<1; measurement protocol outlined in Online Supplemental Resource Fig. 3a, b, e). The persistent enlargement of inner tongues may thus result either from increased membrane growth rate or impaired actin disassembly and myelin membrane compaction(Dillenburg et al, 2018). To address these hypotheses, we first measured the thickness of compacted layers because an increased membrane growth rate would result in a thicker myelin sheath. Plotting myelin thickness against the axon diameters of LCN2*^KD^* mice demonstrated that the myelin thickness was most prominently increased in small axons, thereby implying increased membrane growth in early remyelination (Online Supplemental Resource Fig. 3c, d). We then assessed the expression of MBP, which is required for actin disassembly and myelin membrane compaction(Zuchero et al, 2015). LCN2*^KD^* mice showed significantly increased MBP intensity relative to the scrambled controls (Fig. 3f, g); NF200+ myelin sheaths devoid of MBP were also observed in these mice (Fig. 3h), indicative of noncompact myelin, as NF200 is normally excluded from compact myelin sheaths by MBP(Aggarwal et al, 2013). Finally, we showed that the thicker myelin in LCN2*^KD^* mice was compact (Fig. 2j). Overall, these data indicate that remyelination is increased in the brains of LCN2*^KD^* mice and suggest that high levels of myelin were produced by LCN2-knockdown oligodendrocytes during the recovery phase in the SAH and MS models.

**Fig. 3.**
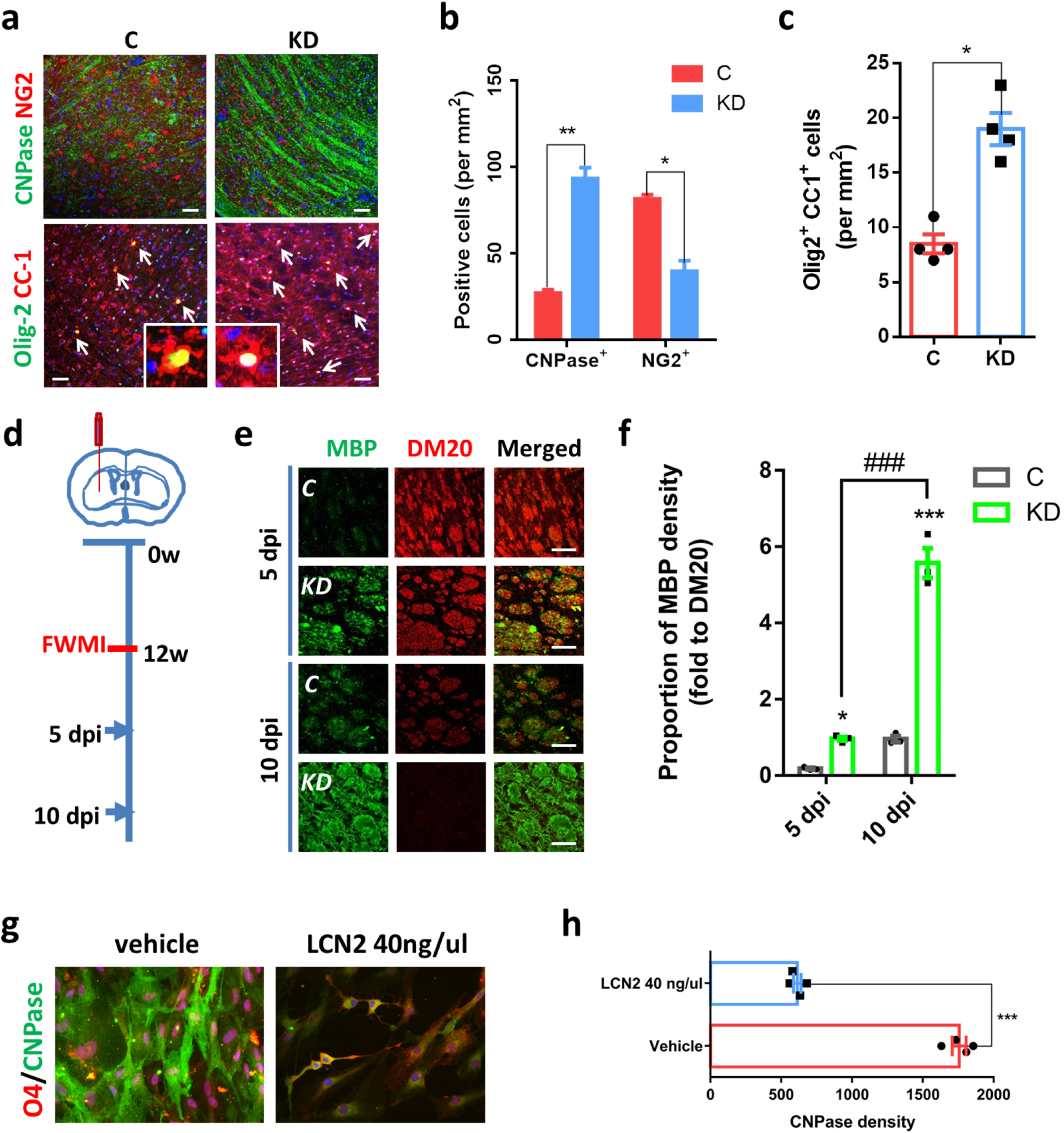
LCN2 inhibits OPC differentiation. **a)** OPC (NG2+), oligodendrocyte (CNPase+), and mature oligodendrocyte lineage (CC1+Olig2+) levels in the white matter of control and LCN2*^KD^* mice at 7 days after SAH. Inset, white arrow indicates CC1+Olig2+ cells. Scale bar, 50 μm. **b)** Cells positive for CNPase and NG2 in control and LCN2*^KD^* mice at 7 days after SAH (n = 4 fields per group). Multiple t test, ***P = 0.0005 (CNPase+), **P = 0.0031 (NG2+). **c)** Number of mature oligodendrocyte lineage cells (CC1+Olig2+) in control and LCN2*^KD^* mice at 7 days after SAH (n = 4 fields per group). Two-tailed Student’s t test, ***P = 0.0004. **d)** Schematic of the timing of FWMI induction. **e)** Distribution of oligodendrocytes (MBP+) and immature oligodendrocytes (DM20+) at 5 and 10 dpi in FWMI. Scale bar, 50 μm. **f)** Relative proportion of the MBP density in control and LCN2*^KD^* mice at 5 and 10 dpi in FWMI (n = 3 fields per group). Two-way ANOVA with Sidak’s multiple comparisons test, *P < 0.05, ***P < vs. the control group, ^###^P < 0.001 vs. 5 dpi. **g)** Image of OPCs (O4+) and oligodendrocytes (CNPase+) in cultured OPCs under vehicle or 40 ng/μLLCN2 treatment with T3 for 24 h. Scale bar, 25 μm. **h)** Analysis of the CNPase fluorescence intensity after vehicle or LCN2 treatment (n = 4 fields per group). Two-tailed Student’s t test, ***P = 0.0008.

### Inhibition of LCN2 does not reduce glial activation and inflammation

Some studies revealed that LCN2 damages the brain by promoting glial activation and proinflammatory factor release(Lee et al, 2015; Nam et al, 2014). Therefore, we asked whether the positive role of LCN2 knockdown in remyelination was attributed to glial activation and inflammatory injury. We evaluated glial activation at 7 days after SAH by immunofluorescence and found no significant reduction in the number of microglia and astrocytes in the white matter between control and LCN2*^KD^* mice (Online Supplemental Resource Fig. 4a-c). Furthermore, we used qPCR to detect the main proinflammatory factors, including IL-1β, TNF-αα, IFN- γ and Ccl2 levels, in the brain tissues. LCN2*^KD^* mice did not show a significant reduction in the mRNA expression level of these factors (Online Supplemental Resource Fig. 4d-g). These data do not support the speculation that the inhibition of LCN2 reduces glial activation and inflammation during remyelination and instead reveal the possibility of an oligodendroglial mechanism amid remyelination(Al Nimer et al, 2016).

**Fig. 4.**
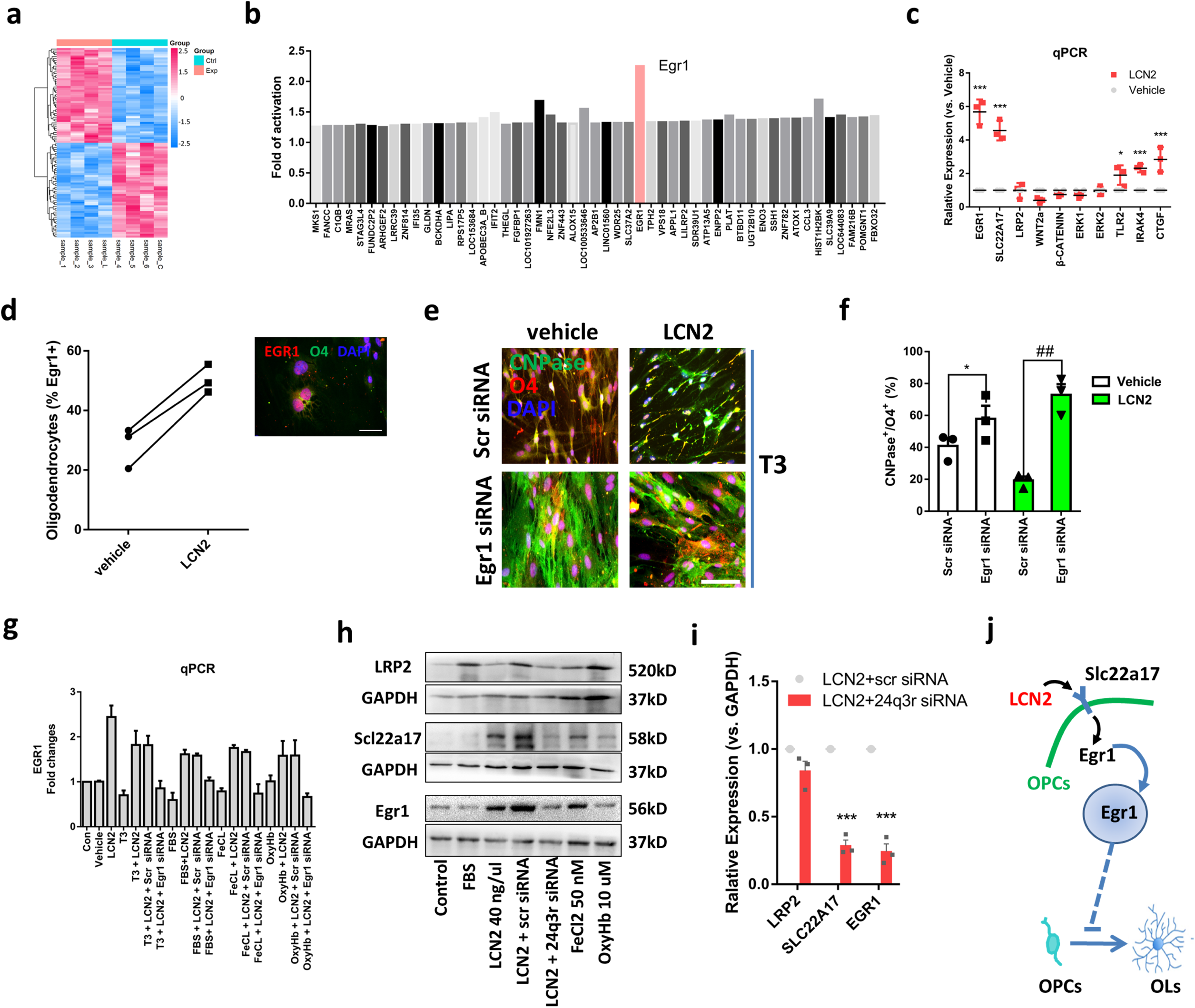
LCN2 activates EGR1 in OPCs. **a)** Clustering map of differentially expressed genes in the vehicle (Ctrl) and LCN2 (Exp) groups. Fold changes in 100 differential expressed genes ranging from -1.5 to 1.5. **b)** Mean fold of activation of some differentially expressed genes in OPCs treated with LCN2 compared with vehicle. EGR1 showed significant activation (Limma moderated t test, P = 0.00015, t = 5.25, df = 6). N = 4 different independent RNA preparations and analyses. The complete list of regulated genes is reported in Supplementary Table 1. **c)** qPCR verification of the relative expression of some differentially expressed genes in OPCs treated with LCN2 compared with vehicle. One-way ANOVA with Dunnett’s’ multiple comparisons test, *P < 0.05, ***P < 0.001 vs. the control group. **d)** Analysis of the EGR1-positive OPCs (O4+) after treatment with vehicle or LCN2. The right panel shows the colocalization of EGR1 and O4 in OPC cultures. Two-tailed Student’s t test, *P = 0.0145. **e)** Image of OPCs (O4+) and oligodendrocytes (CNPase+) in cultured OPCs under scr siRNA or EGR1 siRNA treatment after T3 induction for 24 h. Scale bar, 25 μm. **f)** Proportion of O4+CNPase+ cells in total OPCs under scr siRNA or EGR1 siRNA treatment after T3 induction for 24 h. Two-way ANOVA with Sidak’s multiple comparisons test, *P < 0.05, ^##^P < 0.01 vs. scr siRNA. **g)** qPCR analysis of EGR1-fold changes in cultured OPCs after treatment with vehicle, T3, LCN2, FBS, Fecl2, OxyHb, scr siRNA, EGR1 siRNA, or respective combinations. Two-way ANOVA with Sidak’s multiple comparisons test, *P < 0.05, ^##^P < 0.01. **h)** Western blot of LRP2 (520 kDa), SLC22A17 (58 kDa), EGR1 (56 kDa), and the reference protein GAPDH (37 kDa), after treatment with vehicle, LCN2, FBS, Fecl2, OxyHb, scr siRNA, 24q3r (SLC22A17) siRNA, or their respective combinations. **i)** Relative expression of LRP2, SLC22A17, and EGR1 in OPCs treated with LCN2 plus scr siRNA or 24q3r (SLC22A17) siRNA. Two-way ANOVA with Sidak’s multiple comparisons test, ***P < 0.001. **j)** Schematic diagram of the signaling pathway potentially activated by LCN2 and involved in OPC differentiation. LCN2 increases its membrane receptor SLC22A17 on OPCs and activates the transcription factor EGR1, thereby inhibiting the differentiation of OPCs into oligodendrocytes (OLs).

### LCN2 inhibits OPC differentiation

We next sought to determine whether the cellular mechanism underlying remyelination in LCN2*^KD^* mice involves oligodendroglial responses. At 14 days after SAH, when remyelination occurred, the number of NG2-positive OPCs was decreased in the white matter of LCN2*^KD^* mice compared to the scrambled control (Fig. 3a, b). In contrast, the numbers of oligodendrocytes (CNPase+) were significantly increased (Fig. 3a, b). These data suggest that LCN2 knockdown promotes OPC differentiation into oligodendrocytes after SAH. To specifically address this hypothesis, we analyzed the proportion of Olig2+ CC1+ cells that were mature oligodendrocytes and Olig2+ CC1− cells that were immature cells and found a higher proportion of mature oligodendrocytes in the white matter of LCN2*^KD^* mice (Fig. 3c). Although LCN2 reticence increased the number of Olig2+ CC1+ cells in the white matter after SAH, the total number of Olig2+ cells were not significantly changed, suggesting that LCN2*^KD^* mice show accelerated differentiation into mature oligodendrocytes but no change in OPC proliferation.

In addition to whole brain demyelinating diseases such as SAH and MS, we sought to detect whether the efficient remyelination process is different in a focal WMI model in LCN2*^KD^* mice. Using temporally discrete OPC responses following focal demyelinated lesion induction in white matter (FWMI) (Fig. 3d), the proportions of mature oligodendrocytes (MBP+) and immature oligodendrocytes (DM20+) were assessed at the time of initiation of oligodendrocyte differentiation (5 dpi) and remyelination (10 dpi) (Fig. 3e). LCN2*^KD^* mice had a higher proportion of mature oligodendrocytes (MBP+) after FWMI. This result shows that LCN2 inhibition also improves oligodendrocyte differentiation in FWMI (Fig. 3f).

To directly assess the effects of LCN2 on OPC behavior, we first detected OPC proliferation after SAH induction and LCN2 treatment. After counting the number of OPCs (NG2+Olig2+) at 1, 3, 7, 14, and 28 days post-SAH, we did not find significant OPC proliferation in LCN2*^KD^* mice (Online Supplemental Resource Fig. 5a, b). In addition, a gradient concentration of LCN2 was used to directly treat cultured OPCs, and no significant changes in CCK8 viability were found at most concentrations of LCN2; high concentrations (160 ng/ul) of LCN2 reduced CCK8 viability, probably because of the cytotoxicity of high concentrations (Online Supplemental Resource Fig. 5c). Given that bran injury-induced increases in LCN2 are unlikely to reach such high concentrations(Ranjbar Taklimie et al, 2019), these data exclude an effect of LCN2 on OPC proliferation. We subsequently detected the effect of LCN2 on OPC migration by a cell wound healing assay. After a 24-h observation, the mean migration distance of the 40 ng/µl LCN2 treatment group was not significantly different from that of the null LCN2 treatment group (Online Supplemental Resource Fig. 5d, e). Apart from the high concentration of LCN2, most concentrations of LCN2 did not affect the average migration rate of OPCs compared with the null LCN2 treatment group, suggesting that LCN2 does not affect OPC migration (Online Supplemental Resource Fig. 5f). Finally, we took advantage of T3 to induce OPC differentiation and assessed the effect of LCN2 on this process. Treatment with 40 ng/µL LCN2 abolished the T3-induced differentiation of pre-oligodendrocytes (O4+CNPase-) into oligodendrocytes (O4+CNPase+), hence suggesting the inhibitory role of LCN2 in OPC differentiation (Fig. 3g, h).

**Fig. 5.**
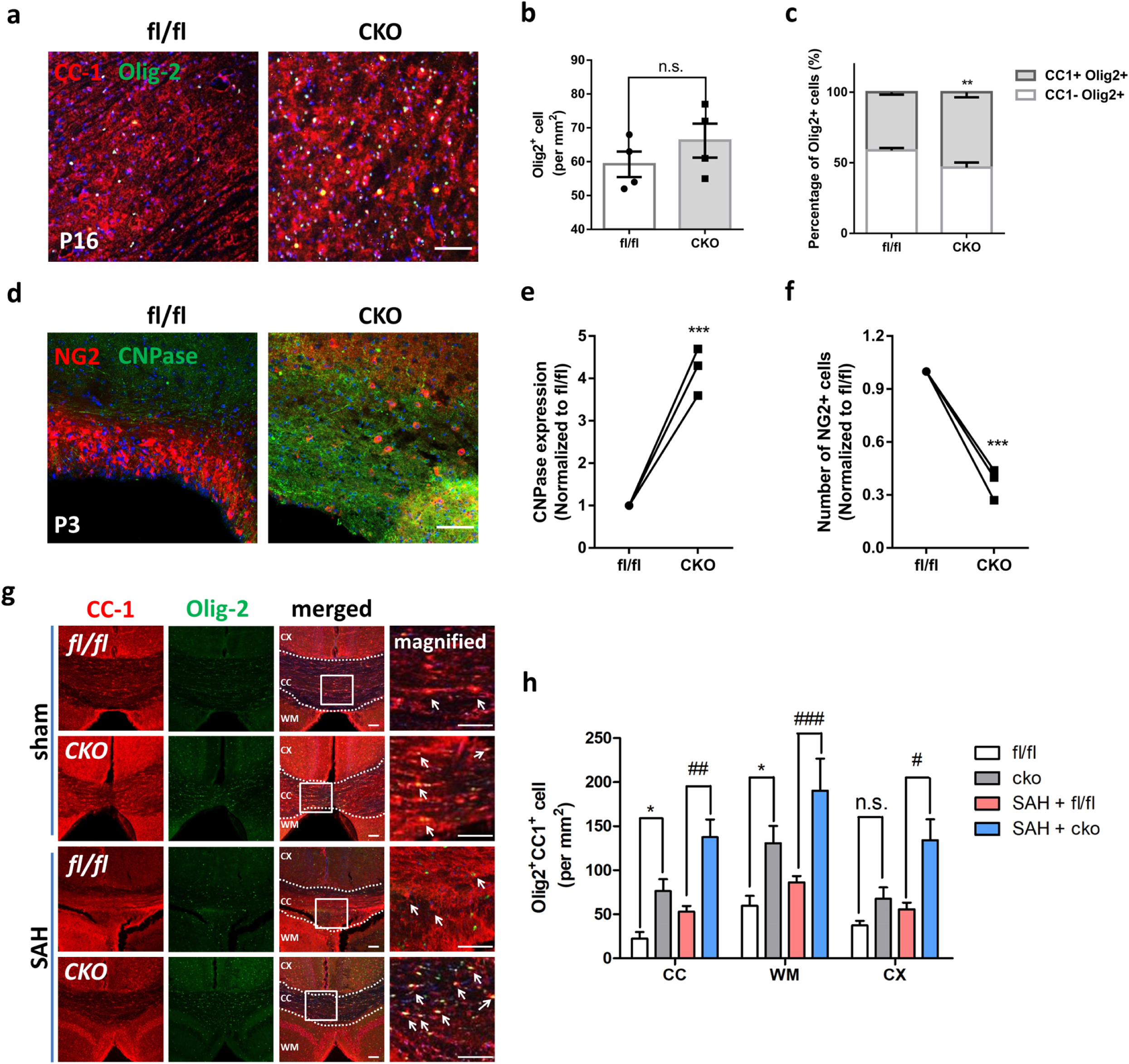
Inactivation of EGR1 promotes OPC differentiation in developing and injured brains. **a)** Image of mature oligodendrocytes (CC1+Olig2+) in fl/fl and CKO mice at P16. Scale bar, 75 μm. **b)** Total number of Olig2+ cells in fl/fl and CKO mice at P16 (n = 4 fields per group). Two-tailed Student’s t test, P = 0.9609. **c)** Percentage of CC1+ and CC1- cells among the total Olig2+ cells in fl/fl and CKO mice at P16 (n = 4 fields per group). Multiple t test, **P = 0.0074. **d)** Image of OPCs (NG2+) and oligodendrocytes (CNPase+) in the subventricular zone of fl/fl and CKO mice at P3. Scale bar, 75 μm. **e)** Normalized CNPase expression of fl/fl and CKO mice at P3. Two-tailed Student’s t test, ***P = 0.0005. **f)** Normalized NG2 expression of fl/fl and CKO mice at P3. Two-tailed Student’s t test, ***P = 0.0002. **g)** Image of mature oligodendrocytes (CC1+Olig2+) in fl/fl and CKO mice at 7 days after sham or SAH, with magnified view of white frame of the corpus callosum (CC). Scale bar, 75 μm. **h)** Analysis of different parts (CC, WM, and CTX) of CC1+Olig2+ cells of fl/fl and CKO mice at 7 days after sham or SAH. Two-way ANOVA with Sidak’s multiple comparisons test, *P < 0.05 vs. sham + fl/fl; ^#^P < 0.05, ^##^P < 0.01, ^###^P < 0.001 vs. SAH + fl/fl.

### LCN2 activates EGR1 in OPCs

The above data suggest that LCN2 is a negative regulatory factor of OPC differentiation after WMI. We next sought to explore the molecular mechanisms underlying OPC differentiation. The activation and inactivation of molecular pathways in LCN2-stimulated oligodendroglial lineage cells for 24-h was simultaneously assessed by measuring the mRNA levels of signaling proteins using an Affymetrix GeneChip Human Gene 1.0ST Array. Differentially expressed genes were normalized to GAPDH expression (The complete list of regulated genes is reported in Supplementary Table 1), and values were then normalized to vehicle control (Fig. 4a). We found that LCN2 reduced the expression of STRBP, DYNC1I2, FMO8P, API5 and ZFN525, but increased the expression of PLAT, NFE2L3, IFIT2, FMN1, and SLC39A9 as verified by RT-PCR (Online Supplemental Resource Fig. 6). Regarding OPC differentiation, LCN2 did not significantly activate the typical lingo-1, Notch, Erb2, PXRγ and histamine receptors 1/3 that modulate oligodendrocyte differentiation. However, LCN2 slightly decreased positive regulatory signaling, including Wnt/β-catenin, sema4G/plexin and ERK1/2, while increasing negative regulatory signaling, such as CTGF and TLR2/IRAK4 (Online Supplemental Resource Fig. 7a). LCN2 did not activate some reported inhibitory factors of oligodendroglial differentiation, but the upregulation in RhoA was ascribed to LCN2-induced stress response (Online Supplemental Resource Fig. 7b). Notably, LCN2 caused a marked activation of the transcription factor early growth response protein 1 (EGR1), which was verified by qPCR and immunostaining (Fig. 4b-d). In fact, increased EGR1-positive OPCs and EGR1 expression were observed after SAH (Online Supplemental Resource Fig. 8a-c). Additionally, LCN2 increased its receptors SLC22A17 but did not enhance LRP2 expression (Fig. 4c). The activation of EGR1 is involved in peripheral myelin development and adult neurogenesis through oligodendrocyte differentiation via neuron-glia signaling(Kwon et al, 2018; Stevens & Fields, 2000). Therefore, the activation of EGR1 may mediate LCN2-regulated remyelination by inhibiting OPC differentiation. To address this deduction, we utilized siRNA to repress the EGR1 activation that accompanied LCN2 treatment. Double-labeling of CNPase and O4 showed that the inhibitory effect of LCN2 on the differentiation of OPCs (O4+CNPase-) into oligodendrocytes (O4+CNPase+) was neutralized in the EGR1 siRNA group compared with the scr siRNA group (Fig. 4e). EGR1 siRNA also promoted T3-induced OPC differentiation (Fig. 4f). Nevertheless, treatment with FBS promoted OPC differentiation, while FeCl2 and OxyHb inhibited OPC differentiation and even reduced EGR1 mRNA levels (Fig. 4g).

**Fig. 6.**
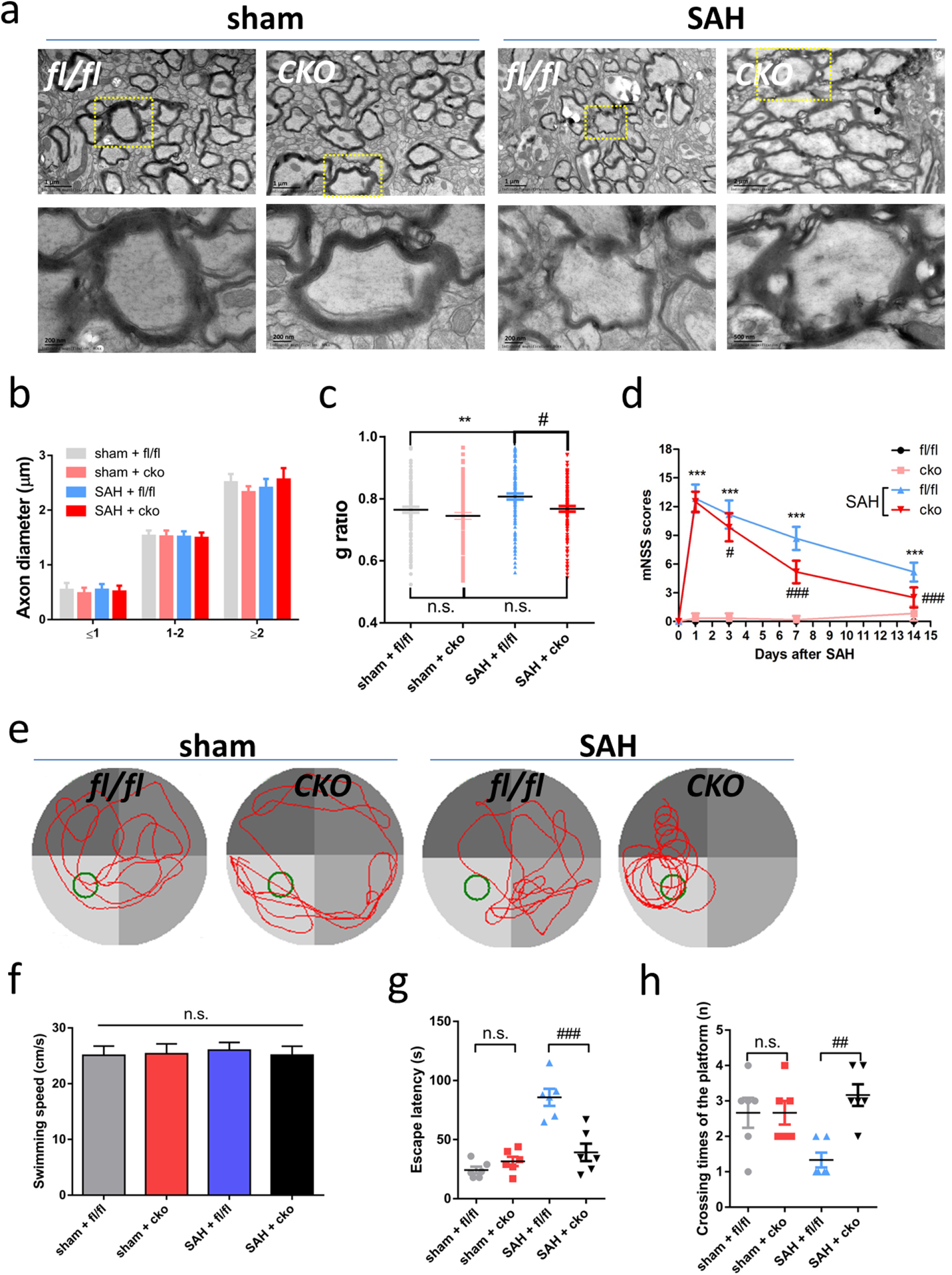
Specific inactivation of EGR1 promotes remyelination and white matter functional recovery. **a)** Ultrastructure of the white matter of fl/fl and CKO mice at 7 days after sham or SAH. The yellow frames are magnified blow. Scale bar, 2 μm or 200 nm. **b)** Axon diameter in fl/fl and CKO mice at 7 days after sham or SAH. Two-way ANOVA with Sidak’s multiple comparisons test, no difference in diameter < 1, 1-2, and < 2. **c)** G-ratio of fl/fl and CKO mice at 7 days after sham or SAH. One-way ANOVA with Dunnett’s’ multiple comparisons test, **P < 0.01, ^#^P < 0.05, vs. SAH + fl/fl. **d)** mNSS evaluation of fl/fl and CKO mice at 0, 1, 3, 7, and 14 days after sham or SAH. Two-way ANOVA with Sidak’s multiple comparisons test, ***P < 0.001 vs. sham + fl/fl, ^#^P < 0.05, ^###^P < 0.001 vs. SAH + fl/fl. **e)** Trajectory of fl/fl and CKO mice in the water maze pool. **f)** Analysis of the swimming speed of fl/fl and CKO mice with sham or SAH. One-way ANOVA with Dunnett’s multiple comparisons test. n.s. indicates no significant. **g)** Escape latency before finding the platform of fl/fl and CKO mice with sham or SAH. One-way ANOVA with Dunnett’s multiple comparisons test. n.s. indicates no significant vs. sham + fl/fl, ^###^P = 0.0002 vs. SAH + fl/fl. **h)** Number of platform crossings within 120 s of fl/fl and CKO mice with sham or SAH. One-way ANOVA with Dunnett’s multiple comparisons test. n.s. indicates no significant vs. sham + fl/fl, ^##^P = 0.0078 vs. SAH + fl/fl.

We next verified whether LCN2 and its receptors regulated EGR1 activation through western blotting. Treatment with LCN2 for 24 h increased SLC22A17 and EGR1 but not LRP2 levels in cultured OPCs (Fig. 4h). This effect of LCN2 on the activation of SLC22A17 and EGR1 was reversed by the administration of 24q3r siRNA (Fig. 4i). FeCl2 showed similar effects on the activation of SLC22A17 and EGR1, suggesting that iron transport may be associated with OPC differentiation(Morath & Mayer-Proschel, 2002). However, OxyHb, which is responsible for iron transport and metabolism such as LCN2, induced no significant changes in the activation of SLC22A17 and EGR1 in OPCs. FBS did not increase the expression of SLC22A17 and EGR1 but regulated upregulated LRP2 expression. Taken together, these data indicate that LCN2 activates SLC22A17/EGR1 signaling but not LRP2, thereby inhibiting the differentiation of OPCs into oligodendrocytes (Fig. 4j).

### Inactivation of EGR1 promotes OPC differentiation in developing and injured brain

EGR1 shows various functional attribute in fibroblasts, epithelial cells, lymphocytes, microvascular endothelial cells and neurons, according to previous studies(Duclot & Kabbaj, 2017). To precisely determine whether EGR1 activation is required for the inhibitory effect of LCN2 on OPC differentiation, we constructed a constitutive conditional knockout in which oligodendroglial lineage cells cannot respond to LCN2 due to EGR1 excision in OPCs (PDGFRa-Cre; EGR1^fl/fl^, CKO) (Online Resource Supplemental Fig. 9a). Tamoxifen effectively reduced the mRNA level of EGR1 in the whole brains of CKO mice (Online Resource Supplemental Fig. 9b). By immunofluorescence, EGR1 expression was confirmed to be eliminated in PDGFRα+ oligodendrocyte lineage cells in tamoxifen-treated CKO mice (Online Resource Supplemental Fig. 9c, d). However, the conditional knockout of EGR1 in OPCs did not affect the number of OPCs (PDGFRα+) in adult mouse brains (Online Resource Supplemental Fig. 9e, f). CKO mice displayed no abnormal behaviors, indicating no obvious myelin damage.

We next assessed whether CKO affects the development of myelination in mice. At P16, when myelination is normally underway, oligodendrocyte lineage cells (Olig2+) were slightly increased in CKO mice; CKO mice had more mature OPCs (CC1+Olig2+) than do fl/fl mice (Fig. 5a-c). The ascendancy of oligodendrocyte maturation in EGR1 conditional knockout mice was observed as early as P3 during development. More oligodendrocytes (CNPase+) were observed in the subventricular zone where OPC proliferation and myelin repair occur (Fig. 5d-f). CKO reduced the number of OPCs (NG2+), suggesting that the inactivation of EGR1 in OPCs facilitates myelination in developing brain. To further analyze whether CKO affects OPC differentiation in pathological scenarios, the CKO mice were subjected to SAH induction (Online Supplemental Resource Fig. 10a). CKO eliminated the increase in EGR1 in OPCs after SAH (Online Supplemental Resource Fig. 10b-c). Then, we used immunofluorescence staining to detect oligodendrocyte maturation at 14 days after SAH. In the sham groups, CKO increased the proportion of mature oligodendrocytes (CC1+Olig2+) in the corpus callosum and white matter. In the SAH groups, the proportion of mature oligodendrocytes (CC1+Olig2+) in the corpus callosum and white matter was increased in CKO mice compared to fl/fl mice (Fig. 5g, h). These data revealed that the inactivation of EGR1 promotes OPC differentiation during development and injury.

### Specific inactivation of EGR1 promotes remyelination and white matter functional recovery

We have shown that the specific inactivation of EGR1 promotes OPC differentiation; thus, the issue of whether EGR1 promotes white matter functional recovery should be answered. At 10 dpi, the number of NG2-positive OPCs located in white matter lesions was decreased in CKO mice compared with fl/fl mice; moreover, the normalized expression of CNPase was significantly increased in CKO mice, indicating that the inactivation of EGR1 promotes remyelination after FWMI (Online Supplemental Resource Fig. 11a-c). During the myelin recovery phage of the MS model (5wpi), the double-labeling results showed that remyelinated MAG expression was increased in the spinal cord of CKO mice; furthermore, a thicker myelin sheath in almost all axon diameters was found in the white matter of CKO mice compared to fl/fl mice (Online Supplemental Resource Fig. 11d-f). To detect the behavior of CKO mice during remyelination, we used BMS scoring and water maze to examine white matter function. Compared with fl/fl mice, the BMS score, which represents the motor function of CKO mice, was increased starting from 4 to 6 wpi (Online Supplemental Resource Fig. 11g). The swimming distance and escape latency were reduced in CKO mice compared with fl/fl mice (Online Supplemental Resource Fig. 11h, i). These data indicate that the inactivation of EGR1 in OPCs promotes remyelination and behavioral recovery in FMWI and MS models.

Previous work revealed that SAH induced obvious demyelination and that remyelination was insufficient even after 14 days of SAH(Kooijman et al, 2014; Li et al, 2018). Remyelination in the ultramicroscopic field was analyzed 14 days after SAH (Fig. 6a). The axon diameter in fl/fl mice was the same as that in CKO mice at all scales (Fig. 6b). In the sham groups, no significant difference in the g-ratio was found between CKO and fl/fl mice. After SAH, the thickness of myelin was increased in the CKO group compared to the fl/fl group (Fig. 6c). In the behavior assessment, CKO did not significantly change the mNSS scores of the sham groups. However, compared to fl/fl mice, CKO mice showed decreased mNSS scores at 3, 7 and 14 days after SAH (Fig. 6d). We next utilized the Morris water maze to detect cognitive repair after SAH (Fig. 6e). There was no significant difference in swimming speed among the sham and SAH groups. In the sham groups, CKO mice showed no significant difference in escape latency, crossing times of the platform and swimming distance from fl/fl mice. In the SAH groups, CKO shortened the swimming distance and escape latency and reduced the number of platform crossings (Fig. 6f-h). In summary, these data indicate that the specific inactivation of EGR1 in OPCs promotes remyelination, thereby providing a promising target for white matter functional recovery.

## DISCUSSION

The identification of underlying mechanisms and new therapeutic targets are urgently required for myelin repair. In this study, we investigated the function of LCN2 and EGR1 in SAH (atypical and acute)- and MS (typical and chronic)-induced WMI diseases. Using a microarray, conditional knockout, in vitro manipulation of oligodendrocyte lineage cells, and ex vivo/in vivo modeling of remyelination, we uncovered the inhibitory effects of LCN2 and activated EGR1 on appropriate oligodendrocyte differentiation in WMI. These findings extend previous studies that did not find an effect of LCN2 on myelin (Chun et al, 2015; Nam et al, 2014), by demonstrating a direct and disease-relevant role in oligodendrocyte lineage cells. Our results also extend the list of negative regulators of oligodendroglial differentiation to now include the regulation of myelin repair and the appropriate response to WMI in human disease(Kremer et al, 2011).

Resident OPCs are the main promoters of remyelination after CNS injury and eventually promote partial clinical remission(Kremer et al, 2016). However, the understanding of efficiency of oligodendrogenesis and remyelination in the adult CNS is limited(Chang et al, 2002) for two complementary reasons. First, OPC reservoirs in the brain appear to be depleted and rarely migrate into lesions, especially during the progressive phase of the disease, but we detected excessive OPCs in the white matter after SAH, consistent with other findings in MS and intracranial hemorrhage(Miron et al, 2011);. Second, and most likely of greater relevance, inhibitors of oligodendroglial differentiation and maturation may act either alone or synergistically to interfere with regeneration processes(Bruce et al, 2010). Inhibitory components, such as LINGO-1, Id2, Id4, IFN-γ, etc., have been identified over the past few years(Kremer et al, 2011). LCN2 was found to be especially present in white matter lesions and to negatively regulate remyelination. Furthermore, we revealed that LCN2 inhibits OPC differentiation, indicating LCN2 activation as a novel oligodendroglial differentiation inhibitor. Although problems with oligodendroglial differentiation are a major reason for remyelination failure, a differentiation blockade may also result from a lack of differentiation-inducing signals(Franklin & Ffrench-Constant, 2008). In the present study, we utilized the thyroid hormone T3, which enhanced oligodendroglial differentiation and proliferation early in development(Franco et al, 2008), to verify the role of LCN2 in the differentiation process. Data showed that LCN2 inhibits T3-induced oligodendroglial differentiation, raising the question of how LCN2 affects T3 signaling. The increased enzymatic deactivation of T3 may play a role in impaired remyelination in experimental autoimmune encephalomyelitis (Castelo-Branco et al, 2014). Thus, whether the T3 signal is still involved in the progress of remyelination should also be addressed in SAH or MS. Overall, these results provide new putative targets for future remyelination therapies.

Numerous experimental and clinical studies have shown that LCN2 is upregulated during acute and chronic injury(Richardson, 2005). Whether LCN2 is protective or injurious after injury, however, is still disputed(Ferreira et al, 2015). The reason is associated with the contentious role of LCN2 in the inflammatory response. Although in hepatology, LCN2-null mice showed a significantly lower recruitment of neutrophils and leukocytes, indicative of protective effects, and LCN2 might act as an intrinsic “help-me” sensor upon injury(Asimakopoulou et al, 2016). In the brain, LCN2 was found to promote neuroinflammation but appears to show neuroprotection in some diseases(Kang et al, 2018). In fact, microarrays showed that LCN2 actually induced an inflammatory response in OPCs when a group of upregulated inflammatory factors was examined after LCN2 stimulation. However, LCN2 reticence is not sufficient to alleviate MS- or SAH-induced neuroinflammation because the inhibition of LCN2 failed to repress the activation of microglia and astrocytes during the two disease scenarios. There might be a parallel mechanism, such as iron dysregulation and neurotoxicity, to induce neuroinflammatory process(Dekens et al, 2018), or very low level of LCN2 could sensitively activate the downstream neuroinflammatory reaction(Bhusal et al, 2019). The precise impact of LCN2 in the CNS is far from completely unraveled and is considered to be multifaceted. From a clinical perspective, LCN2 could be targeted therapeutically to dampen proinflammatory astrocytic activation(Xing et al, 2014). Nonetheless, at this point, the absence of a specific antagonist for LCN2, as well as the poorly elucidated mediation of astrocytic neurotoxicity by LCN2 through binding to specific receptors, makes the task of counteracting LCN2 effects on the progression of CNS diseases very challenging.

The effects of LCN2 on oligodendrocyte differentiation were found to be mediated by the transcription factor EGR1, as its neutralization in OPCs eliminated the LCN2-driven suppression of oligodendrocyte differentiation and hypomyelination. Accordingly, we showed that EGR1 expression in oligodendroglial lineage cells in vivo coincided with oligodendrocyte differentiation and myelin generation during development and following injury. During hypomyelination after injury, there was a concomitant upregulation of EGR1. The concurrent downregulation of several transcription factors, including Id2, Egr1, and Sox11, is critical for OPC differentiation(Swiss et al, 2011), and importantly, the increased expression of these transcription factors was found to impede oligodendrocyte differentiation and myelination during development and after WMI. In esophageal squamous cell carcinoma, EGR1 are the key transcription activation protein of LCN2, within a positive LCN2-MEK/ERK-LCN2 loop, to promote the migration and invasion of esophageal squamous cell carcinoma cells(Zhao et al, 2019). Thus, the downregulation of EGR1 after injury may allow oligodendroglial lineage cells to be more conducive to response and successful remyelination.

The differentiation of OPCs into mature oligodendrocytes has also been associated with the activation or inactivation of other transcription factors, such as PKCα, TCF4, and NFATc4(Kremer et al, 2016; Trimarco et al, 2014). Although we did not detect significant changes in the expression of these genes after LCN2 stimulation in cultured OPCs, in vivo observation in MS and other diseases confirmed the nuclear translocation and activation of oligodendrocyte differentiation-dependent transcription factors, which suggests that complementary pathways likely coordinate oligodendroglial lineage and may also respond in parallel. These LCN2-insensitive and LCN2-sensitive transcription factors may divide into distinct subsets of OPCs, since a single-cell RNAseq database confirmed distinct expression patterns of the abovementioned factors within OPCs(Marques et al, 2016). In addition, different transcription factors may involve distinct cellular regulatory signals. For example, we observed alterative LCN2-induced mechanisms, which have previously been demonstrated to modulate oligodendrocyte differentiation, suggesting that an alternate mechanism may be involved in the LCN2-dependent decrease in oligodendrocyte differentiation during remyelination.

Admittedly, some limitations of the present study should also be mentioned. First, we used an siRNA approach to study the role of LCN2 in remyelination after WMI, yet in view of the efficiency and toxicity of gene interference, mouse models with LCN2 deletion (LCN2^-/-^) or even conditional LCN2 deletion in the brain (e.g., Nestin^cre/cre^/LCN2^fl/fl^) may require further study. Second, recent efforts in high-throughput screening have identified a number of compounds that potently promote OPC differentiation and remyelination(Deshmukh et al, 2013; Kremer et al, 2016); thus, the identification of which pharmacological strategy is feasible and realistic in the regulation of LCN2 or EGR1 is an issue that must be resolved. Third, additional details of how LCN2 modulates EGR1 activation should be addressed through genetic and pharmacological approaches.

In summary, we provide the first evidence fora shared mechanism of LCN2 and EGR1 in both typical/chronic white matter disorders (MS) and atypical/acute white matter disorders (SAH), despite having distinct etiologies. Our data demonstrate the inhibitory properties of LCN2 in OPC differentiation via the activation of the transcription factor EGR1. Thus, we propose that therapies specifically removing LCN2 or inactivating EGR1 in oligodendroglial lineage cells could represent a novel strategy to enhance differentiation and remyelination in WMI patients.

## Materials and methods

### Animals

All animal protocols were approved by the Institutional Animal Management Committee of the Third Military Medical University. All animal studies were conducted in accordance with the China Public Health Service’s Policy on Humane Care and Use of Laboratory Animals. Mouse colonies were maintained in accordance with institutional guidelines. C57BL/6 mouse colonies were established from mice initially purchased from the Animal Center of Third Military Medical University (Permit Number: Yu2017-0002; Chongqing, China). PDGFRα-Cre mice were purchased from the Jackson Laboratory (Cat log: 018280, Bar Harbor, ME). The Egr1^fl/fl^ mice were kindly provided by the Model Animal Research Center of Nanjing University. LCN2^KD^ mice were generated via injection of LCN2 siRNA, which induced stable knockdown in C57BL/6 mice. The mice were housed in an environment with a 12-hour light/dark cycle and allowed free access to food and water throughout the experimental period. To limit the use of animals, power was calculated by a two-sided 95% confidence interval via the normal approximation method using SPSS 13.0 software and reached > 80% power (84–100%) for all experiments. ARRIVE guidelines were followed in providing the details of the experiments and quantifications and in reporting.

### siRNA Administration

According to methods described previously(Zuo et al, 2017), an intracerebroventricular injection was performed. Briefly, anesthetized mice were placed on a stereotaxic apparatus (RWD Life Science, Shenzhen, China), and the bregma point was exposed. A burr hole was drilled into the bone of the left hemisphere, and the coordinates were 1.5 mm lateral, 3.4 mm posterior, and 3.5 mm below the horizontal plane of the bregma. A 2 μL volume of distinct siRNA was delivered into the amygdala with a Hamilton syringe (Hamilton Company, Reno, NV). The siRNAs included 1 μg/μL LCN2 siRNA, EGR1 siRNA, 24q3r (SLC22A17) siRNA or respective scrambled siRNA (scr siRNA) Santa Cruz Biotechnology, Shanghai, China). The injection was performed twice at 12 and 24 h before SAH to enhance the silencing effect.

### Mouse SAH

The endovascular perforation model of SAH in mice was established as reported previously(Chen et al, 2015; Li et al, 2018). Briefly, mice were anesthetized with 5% chloral hydrate (350 mg/kg, intraperitoneal). A sharpened 5-0 monofilament nylon suture was inserted rostrally into the left internal carotid artery from the external carotid artery stump and perforated the bifurcation of the anterior and middle cerebral arteries. Sham-operated mice underwent the same procedures except that the suture was withdrawn without puncture. All animals were kept at 22–25℃,°C and 65–70% humidity, with a 12-h light/dark cycle and were given sufficient food and water. Blood pressure and heart rate were noninvasively monitored during operation via the tail. The severity of SAH was blindly assessed in all animals after they were sacrificed, as previously described(Sugawara et al, 2008). Each animal received a total score by summing the scores. Only SAH mice with moderate hemorrhage (scores of 8-12) were included in the experiments.

### Cuprizone diet-induced MS

Experimental demyelination was induced by feeding male LCN2*^KD^* mice and their respective wild-type littermates a diet containing 0.2% cuprizone (bis-cyclohexanone oxaldihydrazone, Sigma-Aldrich, Shanghai, China) mixed into standard rodent chow(Werneburg et al, 2017). Cuprizone feeding started at postnatal day 56 (P56) and was maintained for 5 weeks to induce profound demyelination. Spontaneous remyelination was enabled by withdrawing cuprizone from the diet. During experimentally induced demyelination and remyelination, the mice were observed daily, and their body weight was measured twice per week.

### Magnetic resonance imaging

Magnetic resonance imaging (MRI) was performed using a 7.0T small animal magnetic resonance system (PharmaScan; Bruker, Ettlingen, Germany). Mice were anesthetized with 1% isoflurane (Shandong Keyuan Pharmaceutical Co. Ltd., Shandong, China), and the heart rates of mice were maintained at ∼100 bpm. At 7 days after SAH or MS withdrawal, T2-weighted turbo spin-echo sequences (T2WI) were collected with the following set of parameters: field of view = × 2cm2 cm 2cm; slice thickness = 1 mm; slices = 15; interslice distance = 1 mm; repetition time = 3 s; averages = 1; matrix size = 256 × 256; flip angle = 180°; and total scan time for image acquisition =1 min 20s. Five reference images were obtained at 14 days with the following set of parameters: field of view = 2 cm; slice thickness = 0.6 mm; slices = 20; matrix size = 128 × 128; repetition time = 5 s; and average = 2. The WMI volume was calculated with ImageJ software (National Institutes of Health, Bethesda, MD) using the following equation: injury size % =100 × Σ{injury area−(ipsilateral hemisphere−contralateral hemisphere)}/Σ(contralateral hemisphere). Fractional anisotropy (FA) was measured from the tensor map in the ipsilateral internal capsule (IC) with ParaVision 5.0 software (Bruker, Ettlingen, Germany).

### Neurological function evaluation

*mNSS*. As previously reported(Xu et al, 2017), neurological functions were evaluated by a modified neurological severity score (mNSS) method at 1, 3, 5, 7, 9, 12, and 14 days after SAH. The mean of the neurological scores determined by two blinded observers was calculated.

*BMS*. Recovery of hindlimb motor function was scored by the Basso mouse scale (BMS) open-field locomotor rating scale, which was developed specifically for mice(Basso et al, 2006). The score ranges from 0 (complete paralysis) to 9 (normal mobility), and the number of errors at each footstep was measured for 100 steps. Mice were observed individually for 2 min in an open field.

*Morris Water Maze*. The Morris water maze was performed to evaluate hippocampus-dependent spatial learning and memory in mice(Lee et al, 2019). A large circular tank (0.8 m diameter, 0.4 m depth) was filled with water (25 ± 1°C, 20 cm depth), and the escape platform (8×4 cm) was submerged 1 cm below the surface. Each section was monitored by a video capture system. The escape latency and trajectory of swimming were recorded for each mouse.

The hidden platform was located at the center of one of the four quadrants in the tank. The location of the platform was fixed throughout testing. Mice had to navigate using extramaze cues that were placed on the walls of the maze. From days 1 to 4, mice underwent three trials with an intertrial interval of 5 min. The mice were randomly placed in the tank facing the sidewall at one of the four start locations and allowed to swim until they found the platform or for a maximum of 120 s. Each mouse that failed to find the platform within 120 s was guided to the platform. The animal then remained on the platform for 20 s before being removed from the pool. The day after the hidden platform training, a probe trial was conducted to determine whether mice used a spatial strategy to find the platform. On day 5, the platform was removed from the pool, and the mouse was allowed to swim freely for 120 s. The time spent (escape latency), swimming distance, and the number of times the mice crossed the former position of the hidden platform in each quadrant of the pool were recorded.

### Black Gold II myelin staining

The Black Gold II staining kit (Merck Millipore, Chengdu, China) was used according to the manufacturer’s instructions(Santiago Gonzalez et al, 2017). Briefly, 50-μm paraformaldehyde- (PFA)fixed brain sections were mounted onto Superfrost Plus slides (Fisher Scientific, Shanghai, China). Coronal brain slices were initially air dried and then rehydrated and transferred to a lukewarm 0.3% Black Gold II solution. After color development (10 min), the slides were rinsed with a 1% sodium thiosulfate solution at 60°C, dehydrated, and mounted with Permount. The integrated staining intensity in several brain areas was assessed by ImageJ software (National Institutes of Health, Bethesda, MD). Data represent the pooled results from at least four brains per experimental group. Ten slices per brain were used, and quantification was performed by researchers blinded to the genotype of the sample.

### Transmission electron microscope analysis

Mice were intracardially perfused with 4% (w/v) PFA and 2% (v/v) glutaraldehyde (Aladdin Industrial Corporation, Shanghai, China) in 0.1 M phosphate buffer. Tissue was postfixed overnight at 4°C and transferred to 1% (v/v) glutaraldehyde until embedding. Tissue sections (1 mm) were processed into araldite resin blocks. Additionally, 1-μm microtome-cut sections were stained with a 1% toluidine blue/2% sodium borate solution prior to bright field imaging at 100× magnification using a Zeiss AX10 microscope. The number of myelinated axons was blindly quantified in 50 μm × 50 μm images of corpus callosum, with 2–4 sections counted per mouse and then the values were averaged. Ultrathin sections (60 nm) were cut from the corpus striatum and stained in uranyl acetate and lead citrate, and the grids were imaged on a JEOL transmission electron microscope. Axon diameter, myelin thickness, and inner tongue thickness were calculated from the measured area based on the assumption of circularity using ImageJ software 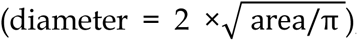, and a minimum of 100 axons per animal were analyzed. G-ratios (the ratio of the axon diameter to the axon plus myelin sheath diameter) were calculated using ImageJ software for at least 100 axons per animal. The inner tongue thickness was calculated by subtracting the axon diameter from the diameter of the innermost compact layer.

### FWMI

Focal white matter demyelinating lesions were induced in the cerebral striatum of 12-week-old male C57BL/6 mice by stereotaxic injection of 4 μL of 0.01% (v/v) ethidium bromide (Sigma-Aldrich, Shanghai, China) using a Hamilton syringe. At 5 and 10 days after injection (dpi), mice were intracardially perfused with 4% PFA, cryoprotected, cryosectioned, and stained as above. Nonlesioned focal white matter injury (FWMI) served as a control.

### OPC cultures

Mixed glial cultures of OPCs were generated from both female and male C57BL/6 P0–P2 mouse pups in vitro, as reported(Weil et al, 2019). The mixed cultures were placed on a rotary shaker at 37°C at 250 rpm for 1 h to de-adhere microglia. OPCs were subsequently isolated from the floating fraction after a 16-h incubation on a rotary shaker, and the contaminating astrocytes were depleted by differential adhesion. OPCs were plated in DMEM/F12 containing 50 μg/ml apo-transferrin, 30 nM sodium selenite, 0.1%(w/v) BSA, 60 ng/ml progesterone, 16 μg/ml Putrescine, 10 μg/ml insulin, 100 μg/ml bovine serum albumin fraction V, 400 ng/ml L-thyroxine, 4.5 g/L glucose, 1% (v/v) L-glutamine, 1% (v/v) pyruvate, 1% penicillin/streptomycin, 10 ng/ml platelet-derived growth factor, and 10 ng/ml fibroblast growth factor-2 (all from Sigma-Aldrich, Shanghai, China). OPCs were planted at 2 × 10^4^ cells per well in 6-well polylysine (0.1 μg/ml)-coated plates (Corning Incorporated, Coring, NY) or confocal plate (glass bottom dish, Cellvis, Mountain View, CA). OPCs were treated with LCN2 (1, 2.5, 5, 10, 20, 40, 80, or 160 ng/ml, R&D Systems, Shanghai, China) or vehicle control (0.0002% BSA) for 1 day. In a subset of experiments, OPCs were treated with 400 ng/ml triiodothyroxine (T3) and LCN2 (40 ng/ml) or vehicle, or plus 5 ng/ml EGR1 siRNA and scr siRNA control. In the molecular mechanism experiments, OPCs were treated with 5% (v/v) FBS, 10 μM oxyhemoglobin (OxyHb), 50 nM Fecl2, 5 ng/ml 24q3r (SLC22A17) siRNA and scr siRNA control, and their respective combinations. Cells were matured to oligodendrocytes by withdrawing growth factors from the media. For CNPase intensity measurements, O4 and CNPase staining was measured in square pixels (px2) with a threshold of 2000 px2 set to exclude background/false positives. An average of 100 cells was counted per image, with 2 images assessed per condition per biological replicate (a total of > 600 cells quantified per condition); values were averaged per biological sample.

### Microarray analysis

Human OPCs (ScienCell Research Laboratories, Carlsbad, CA) were grown in Oligodendrocyte Precursor Cell Medium (OPCM, ScienCell Research Laboratories, Carlsbad, CA), treated with LCN2 or vehicle for 24 h, and harvested. Total RNA was extracted with Trizol (Invitrogen, Shanghai, China). RNA was further purified, and genomic DNA was eliminated using the RNEasy Plus mini kit (Qiagen, Shanghai, China). Human GeneChip® Whole Transcript Expression microarrays were obtained from Affymetrix®. Four independent biological replicates were assayed per group. Sample labeling and array hybridization were conducted by Shanghai Qimin Biotech according to Affymetrix procedures. Data were analyzed with Partek GS using the Affymetrix extended probeset annotation with a GC background-correction RMA algorithm. Differentially expressed mRNAs and probesets were identified using a two-sample t test, with an FDR of 0.1, unless otherwise noted.

### qPCR

Total RNA was extracted from cultured OPCs or brain tissues using the technique described previously by our group(Li et al, 2016). Afterwards, cDNA was synthesized using the High-Capacity cDNA Archive Kit (Qiagen, Shanghai, China). For qualitative reverse transcription PCR (qPCR), 2 ng of cDNA was used, and for analysis, the CFX96 Touch Real-Time PCR Detection System (Bio-Rad, Shanghai, China) was used under the following conditions: 95°C for 20 s, 95°C, for 1 s and60°C for 20 s for 45 cycles (values above 40 cycles were defined as not expressed). All genes were run in triplicate, and each run contained the reference gene (GAPDH) as an internal control to normalize the expression of the target genes. The qPCR assay primers are listed in the Major Resources Tables. The ΔCT of ’normal’ controls was subtracted from the ΔCT of the LCN2-treated group to determine the differences (ΔΔCT) and fold change (2^-ΔΔCT) in gene expression. Gene expression was illustrated by the log10 of fold change values compared to controls.

### Genotyping

Genomic DNA was extracted from tail tissue using the Wizard SV genomic purification system (Promega Biotech, Beijing, China) according to the manufacturer’s instructions. EGR1 floxed mice were genotyped using PCR strategies as previously described(Dillenburg et al, 2018). Briefly, EGR1 floxed mice were genotyped using the primers P1 (CCT TTC CTC ACT CAC CCA CCA TGG) and P2 (CAC CCA CGC AGC TTG AGT TCT C). PDGFRa-Cre mice were genotyped using the primers F (TCA GCC TTA AGC TGG GAC AT), and R (ATG TTT AGC TGG CCC AAA TG). Cre-mediated recombination was detected using P4 (CAA ATG TTG CTT GTC TGG TG) and P5 (GTC AGT CGA GTG CAC AGT TT).

### Fluorescent immunohistochemistry

Brain tissue or spinal cord sections (10-μm-thick) and cultured OPCs were stained using fluorescent immunohistochemistry. Sections or confocal plates were fixed with 4% PFA, and membrane proteins were solubilized with 0.3% Triton X-100 for 30 min. Antigen retrieval, if necessary, was performed by treatment with 95% formic acid for 5 min, followed by boiling in citrate buffer (10 mM sodium citrate, 0.05% Tween-20; pH 6). Sections or confocal plates were blocked with 10% normal goat serum and incubated overnight at 4°C with a combination of antibodies. All primary antibodies were diluted in 4% normal goat serum in PBS. Sections or confocal plates were then incubated for 2 h at room temperature with appropriate fluorescent secondary antibodies. Sections or confocal plates were counterstained with DAPI (Lineage Cell Therapeutics, Alameda, CA) and coverslipped (PermaFluor, Thermo Fisher Scientific, Shanghai, China). To measure the intensity of myelin basic protein (MBP) expression, MBP staining was measured in square pixels (px2) with a threshold of 2000 px2 set to exclude background/false positives.

### Western blotting

Cultured OPCs or brain samples were lysed with RIPA buffer (Thermo Fisher Scientific, Shanghai, China) supplemented with 1% protease inhibitor cocktail set III ethylenediaminetetraacetic acid(EDTA)-free (Roche, Shanghai, China). Protein concentrations were determined using a Pierce BCA Protein Assay Kit (Lineage Cell Therapeutics, Alameda, CA) according to the manufacturer’s instructions. Samples were diluted in loading buffer (Lineage Cell Therapeutics, Alameda, CA) and 5% β-mercaptoethanol (Aladdin Industrial Corporation, Shanghai, China) and heated at 100°C for 5 min, and 10 μg of protein was loaded onto an acrylamide gel (6–15%; Bio-Rad). Gel electrophoresis was performed in Tris- glycine-sodium dodecyl sulfate (SDS) running buffer (Lineage Cell Therapeutics, Alameda, CA) at 100 V, and proteins were transferred onto polyvinylidene difluoride (PVDF) membranes (Merck Millipore, Chengdu, China) for 2 h at 150 mA in 10% transfer buffer [3% Tris–HCl (Aladdin Industrial Corporation, Shanghai, China), 15% glycine (Solarbio Life Sciences, Beijing, China), pH 8.3] and 20% methanol (Shanghai Songon, Shanghai, China) diluted in water. Membranes were blocked with 5% bovine serum albumin in Tris-buffered saline (TBST) [4% sodium chloride (NaCl), 0.1% potassium chloride (KCl), 1.5% Tris–HCl, and 0.1% Tween-20 (all from Aladdin Industrial Corporation, Shanghai, China), pH 7.4] for 1 h at room temperature on a horizontal shaker and then incubated overnight at 4°C with primary antibodies. The membranes were washed three times in TBST for 5 min and incubated with horseradish peroxidase (HRP)-IgG secondary antibody conjugates (1:2000; ZSGB-Bio, Beijing, China) for 1 h at room temperature. A chemiluminescent substrate detection reagent, WesternBright Quantum HRP substrate (Advansta, Menlo Park, CA), was used to visualize bands. For loading control purposes, all membranes were reblotted with anti-mouse or anti-rabbit GAPDH.

### Statistical analyses

Data handling and statistical processing were performed using Microsoft Excel and GraphPad Prism 6.0 software. Data are expressed as the mean ± SEM. Power calculations for sample size were performed (using OpenEpi) and showed power between 84 and 100% for all experiments. All cell counts and analyses were performed blind by researchers blinded to the experimental treatment. Statistical tests included a one-sample t test for data where the values were normalized to control, and a two-tailed Student’s t test when only two groups. were compared Multiple t test was performed when comparing between 2 independent groups. Nonparametric one-way ANOVA with Dunn’s multiple comparison post hoc test was performed when > 3 comparisons were made, and one-way ANOVA with Bonferroni’s multiple comparison post hoc test when was performed for≤ 3 comparisons. Two-way ANOVA with Sidak’s multiple comparisons test was used for intergroup comparison between grouped data. The slopes of myelin thickness versus axon diameter were compared using the Extra Sum of Squares F test. Curve distributions of the proportion of myelinated axons per axon diameter were compared using the Kolmogorov–Smirnov test. P < 0.05 was considered to be statistically significant.

## Acknowledgements

This study was funded by the Top-notch Talent Cultivation Plan of Southwest Hospital (SWH2018BJKJ-05) and the National Natural Science Foundation of China (81220108009 and 81901216)

We thank the Model Animal of Research Center of Nanjing University for providing EGR1^fl/fl^ mice. MS tissue was kindly provided by Dr. Wang Fei from the Third Military Medical University. mRNA microarray scanning and analyze service was provided by Gene-Cloud of Biotechnology Information (Shanghai Qiming Information Technology Co., Ltd.). We also thank Bin Liao, Jinyu Wang, Jie Wang for technical assistance. We thank Prof. Xin Liu, and Prof. Haiwei Xu for helpful discussions.

## Author Contributions

FH and CYJ conceived the project; LQ, RXF, ZHL, QJ, PPY, JHZ, RHZ, and LCJ designed and performed experiments, acquired and analyzed data; RXF performed focal in vivo lesioning; LQ assisted in image analysis protocols; LQ and RXF generated the CKO mice; ZHL and QJ performed neuropathological selection of perinatal brain tissue; PPY performed the neural behavior analysis; LQ, CYJ, and RXF made the figures; LQ wrote the manuscript; CYJ, FH edited the manuscript. All authors read and approved the manuscript.

## References

Aggarwal S, Snaidero N, Pahler G, Frey S, Sanchez P, Zweckstetter M, Janshoff A, Schneider A, Weil MT, Schaap IA et al (2013) Myelin membrane assembly is driven by a phase transition of myelin basic proteins into a cohesive protein meshwork. PLoS biology 11: e1001577

Al Nimer F, Elliott C, Bergman J, Khademi M, Dring AM, Aeinehband S, Bergenheim T, Romme Christensen J, Sellebjerg F, Svenningsson A et al (2016) Lipocalin-2 is increased in progressive multiple sclerosis and inhibits remyelination. Neurology(R) neuroimmunology & neuroinflammation 3: e191

Asimakopoulou A, Borkham-Kamphorst E, Tacke F, Weiskirchen R (2016) Lipocalin-2 (NGAL/LCN2), a “help-me” signal in organ inflammation. Hepatology 63: 669–671

Basso DM, Fisher LC, Anderson AJ, Jakeman LB, McTigue DM, Popovich PG (2006) Basso Mouse Scale for locomotion detects differences in recovery after spinal cord injury in five common mouse strains. Journal of neurotrauma 23: 635–659

Bhusal A, Rahman MH, Lee WH, Bae YC, Lee IK, Suk K (2019) Paradoxical role of lipocalin-2 in metabolic disorders and neurological complications. Biochem Pharmacol 169: 113626

Bruce CC, Zhao C, Franklin RJ (2010) Remyelination - An effective means of neuroprotection. Hormones and behavior 57: 56–62

Castelo-Branco G, Stridh P, Guerreiro-Cacais AO, Adzemovic MZ, Falcao AM, Marta M, Berglund R, Gillett A, Hamza KH, Lassmann H et al (2014) Acute treatment with valproic acid and l-thyroxine ameliorates clinical signs of experimental autoimmune encephalomyelitis and prevents brain pathology in DA rats. Neurobiology of disease 71: 220–233

Chang A, Tourtellotte WW, Rudick R, Trapp BD (2002) Premyelinating oligodendrocytes in chronic lesions of multiple sclerosis. The New England journal of medicine 346: 165–173

Chen Y, Zhang Y, Tang J, Liu F, Hu Q, Luo C, Tang J, Feng H, Zhang JH (2015) Norrin protected blood-brain barrier via frizzled-4/beta-catenin pathway after subarachnoid hemorrhage in rats. Stroke 46: 529–536

Chun BY, Kim JH, Nam Y, Huh MI, Han S, Suk K (2015) Pathological Involvement of Astrocyte-Derived Lipocalin-2 in the Demyelinating Optic Neuritis. Investigative ophthalmology & visual science 56: 3691–3698

Dekens DW, Naude PJW, Keijser JN, Boerema AS, De Deyn PP, Eisel ULM (2018) Lipocalin 2 contributes to brain iron dysregulation but does not affect cognition, plaque load, and glial activation in the J20 Alzheimer mouse model. Journal of neuroinflammation 15: 330

Deshmukh VA, Tardif V, Lyssiotis CA, Green CC, Kerman B, Kim HJ, Padmanabhan K, Swoboda JG, Ahmad I, Kondo T et al (2013) A regenerative approach to the treatment of multiple sclerosis. Nature 502: 327–332

Dillenburg A, Ireland G, Holloway RK, Davies CL, Evans FL, Swire M, Bechler ME, Soong D, Yuen TJ, Su GH et al (2018) Activin receptors regulate the oligodendrocyte lineage in health and disease. Acta neuropathologica 135: 887–906

Duclot F, Kabbaj M (2017) The Role of Early Growth Response 1 (EGR1) in Brain Plasticity and Neuropsychiatric Disorders. Frontiers in behavioral neuroscience 11: 35

Egashira Y, Hua Y, Keep RF, Xi G (2014) Acute white matter injury after experimental subarachnoid hemorrhage: potential role of lipocalin 2. Stroke 45: 2141–2143

Egashira Y, Zhao H, Hua Y, Keep RF, Xi G (2015) White Matter Injury After Subarachnoid Hemorrhage: Role of Blood-Brain Barrier Disruption and Matrix Metalloproteinase-9. Stroke 46: 2909–2915

Ferreira AC, Da Mesquita S, Sousa JC, Correia-Neves M, Sousa N, Palha JA, Marques F (2015) From the periphery to the brain: Lipocalin-2, a friend or foe? Progress in neurobiology 131: 120–136

Franco PG, Silvestroff L, Soto EF, Pasquini JM (2008) Thyroid hormones promote differentiation of oligodendrocyte progenitor cells and improve remyelination after cuprizone-induced demyelination. Experimental neurology 212: 458–467

Franklin RJ, Ffrench-Constant C (2008) Remyelination in the CNS: from biology to therapy. Nature reviews Neuroscience 9: 839–855

Kang SS, Ren Y, Liu CC, Kurti A, Baker KE, Bu G, Asmann Y, Fryer JD (2018) Lipocalin-2 protects the brain during inflammatory conditions. Molecular psychiatry 23: 344–350

Khalil M, Renner A, Langkammer C, Enzinger C, Ropele S, Stojakovic T, Scharnagl H, Bachmaier G, Pichler A, Archelos JJ et al (2016) Cerebrospinal fluid lipocalin 2 in patients with clinically isolated syndromes and early multiple sclerosis. Multiple sclerosis 22: 1560–1568

Kooijman E, Nijboer CH, van Velthoven CT, Mol W, Dijkhuizen RM, Kesecioglu J, Heijnen CJ (2014) Long-term functional consequences and ongoing cerebral inflammation after subarachnoid hemorrhage in the rat. PloS one 9: e90584

Kremer D, Aktas O, Hartung HP, Kury P (2011) The complex world of oligodendroglial differentiation inhibitors. Annals of neurology 69: 602–618

Kremer D, Gottle P, Hartung HP, Kury P (2016) Pushing Forward: Remyelination as the New Frontier in CNS Diseases. Trends in neurosciences 39: 246–263

Kwon W, Kim HS, Jeong J, Sung Y, Choi M, Park S, Lee J, Jang S, Kim SH, Lee S et al (2018) Tet1 overexpression leads to anxiety-like behavior and enhanced fear memories via the activation of calcium-dependent cascade through Egr1 expression in mice. FASEB journal : official publication of the Federation of American Societies for Experimental Biology 32: 390–403

Lee HT, Lee KI, Lin HC, Lee TS (2019) Genetic Deletion of Soluble Epoxide Hydroxylase Causes Anxiety-Like Behaviors in Mice. Molecular neurobiology 56: 2495–2507

Lee S, Jha MK, Suk K (2015) Lipocalin-2 in the Inflammatory Activation of Brain Astrocytes. Critical reviews in immunology 35: 77–84

Li Q, Chen Y, Li B, Luo C, Zuo S, Liu X, Zhang JH, Ruan H, Feng H (2016) Hemoglobin induced NO/cGMP suppression Deteriorate Microcirculation via Pericyte Phenotype Transformation after Subarachnoid Hemorrhage in Rats. Scientific reports 6: 22070

Li Q, Zhao H, Pan P, Ru X, Zuo S, Qu J, Liao B, Chen Y, Ruan H, Feng H (2018) Nexilin Regulates Oligodendrocyte Progenitor Cell Migration and Remyelination and Is Negatively Regulated by Protease-Activated Receptor 1/Ras-Proximate-1 Signaling Following Subarachnoid Hemorrhage. Frontiers in neurology 9: 282

Macdonald RL, Schweizer TA (2017) Spontaneous subarachnoid haemorrhage. Lancet 389: 655–666

Marques S, Zeisel A, Codeluppi S, van Bruggen D, Mendanha Falcao A, Xiao L, Li H, Haring M, Hochgerner H, Romanov RA et al (2016) Oligodendrocyte heterogeneity in the mouse juvenile and adult central nervous system. Science 352: 1326–1329

Miron VE, Kuhlmann T, Antel JP (2011) Cells of the oligodendroglial lineage, myelination, and remyelination. Biochimica et biophysica acta 1812: 184–193

Morath DJ, Mayer-Proschel M (2002) Iron deficiency during embryogenesis and consequences for oligodendrocyte generation in vivo. Developmental neuroscience 24: 197–207

Nam Y, Kim JH, Seo M, Kim JH, Jin M, Jeon S, Seo JW, Lee WH, Bing SJ, Jee Y et al (2014) Lipocalin-2 protein deficiency ameliorates experimental autoimmune encephalomyelitis: the pathogenic role of lipocalin-2 in the central nervous system and peripheral lymphoid tissues. The Journal of biological chemistry 289: 16773–16789

Ngo ST, Steyn FJ, Huang L, Mantovani S, Pfluger CM, Woodruff TM, O’Sullivan JD, Henderson RD, McCombe PA (2015) Altered expression of metabolic proteins and adipokines in patients with amyotrophic lateral sclerosis. Journal of the neurological sciences 357: 22–27

Ranjbar Taklimie F, Gasterich N, Scheld M, Weiskirchen R, Beyer C, Clarner T, Zendedel A (2019) Hypoxia Induces Astrocyte-Derived Lipocalin-2 in Ischemic Stroke. International journal of molecular sciences 20

Rathore KI, Berard JL, Redensek A, Chierzi S, Lopez-Vales R, Santos M, Akira S, David S (2011) Lipocalin 2 plays an immunomodulatory role and has detrimental effects after spinal cord injury. The Journal of neuroscience : the official journal of the Society for Neuroscience 31: 13412–13419

Richardson DR (2005) 24p3 and its receptor: dawn of a new iron age? Cell 123: 1175–1177

Santiago Gonzalez DA, Cheli VT, Zamora NN, Lama TN, Spreuer V, Murphy GG, Paez PM (2017) Conditional Deletion of the L-Type Calcium Channel Cav1.2 in NG2-Positive Cells Impairs Remyelination in Mice. The Journal of neuroscience : the official journal of the Society for Neuroscience 37: 10038–10051

Stevens B, Fields RD (2000) Response of Schwann cells to action potentials in development. Science 287: 2267–2271

Sugawara T, Ayer R, Jadhav V, Zhang JH (2008) A new grading system evaluating bleeding scale in filament perforation subarachnoid hemorrhage rat model. Journal of neuroscience methods 167: 327–334

Swiss VA, Nguyen T, Dugas J, Ibrahim A, Barres B, Androulakis IP, Casaccia P (2011) Identification of a gene regulatory network necessary for the initiation of oligodendrocyte differentiation. PloS one 6: e18088

Trimarco A, Forese MG, Alfieri V, Lucente A, Brambilla P, Dina G, Pieragostino D, Sacchetta P, Urade Y, Boizet-Bonhoure B et al (2014) Prostaglandin D2 synthase/GPR44: a signaling axis in PNS myelination. Nature neuroscience 17: 1682–1692

van Tilborg E, de Theije CGM, van Hal M, Wagenaar N, de Vries LS, Benders MJ, Rowitch DH, Nijboer CH (2018) Origin and dynamics of oligodendrocytes in the developing brain: Implications for perinatal white matter injury. Glia 66: 221–238

Weil MT, Schulz-Eberlin G, Mukherjee C, Kuo-Elsner WP, Schafer I, Muller C, Simons M (2019) Isolation and Culture of Oligodendrocytes. Methods in molecular biology 1936: 79–95

Werneburg S, Fuchs HLS, Albers I, Burkhardt H, Gudi V, Skripuletz T, Stangel M, Gerardy-Schahn R, Hildebrandt H (2017) Polysialylation at Early Stages of Oligodendrocyte Differentiation Promotes Myelin Repair. The Journal of neuroscience : the official journal of the Society for Neuroscience 37: 8131–8141

Wu Y, Peng J, Pang J, Sun X, Jiang Y (2017) Potential mechanisms of white matter injury in the acute phase of experimental subarachnoid haemorrhage. Brain 140: e36

Xing C, Wang X, Cheng C, Montaner J, Mandeville E, Leung W, van Leyen K, Lok J, Wang X, Lo EH (2014) Neuronal production of lipocalin-2 as a help-me signal for glial activation. Stroke 45: 2085–2092

Xu X, Gao W, Cheng S, Yin D, Li F, Wu Y, Sun D, Zhou S, Wang D, Zhang Y et al (2017) Anti-inflammatory and immunomodulatory mechanisms of atorvastatin in a murine model of traumatic brain injury. Journal of neuroinflammation 14: 167

Zhao Y, Xia Q, Liu Y, Bai W, Yao Y, Ding J, Lin L, Xu Z, Cai Z, Wang S et al (2019) TCF7L2 and EGR1 synergistic activation of transcription of LCN2 via an ERK1/2-dependent pathway in esophageal squamous cell carcinoma cells. Cell Signal 55: 8–16

Zuchero JB, Fu MM, Sloan SA, Ibrahim A, Olson A, Zaremba A, Dugas JC, Wienbar S, Caprariello AV, Kantor C et al (2015) CNS myelin wrapping is driven by actin disassembly. Developmental cell 34: 152–167

Zuo S, Ge H, Li Q, Zhang X, Hu R, Hu S, Liu X, Zhang JH, Chen Y, Feng H (2017) Artesunate Protected Blood-Brain Barrier via Sphingosine 1 Phosphate Receptor 1/Phosphatidylinositol 3 Kinase Pathway After Subarachnoid Hemorrhage in Rats. Molecular neurobiology 54: 1213–1228

